# Unveiling the ecology, taxonomy and metabolic capabilities of MBA03, a potential key player in anaerobic digestion

**DOI:** 10.1101/2023.09.08.556800

**Authors:** Roser Puchol-Royo, Javier Pascual, Asier Ortega-Legarreta, Pascal Otto, Jeroen Tideman, Sjoerd-Jan de Vries, Christian Abendroth, Kristie Tanner, Manuel Porcar, Adriel Latorre-Perez

**Affiliations:** Darwin Bioprospecting Excellence, S.L. Parc Cientific Universitat de Valencia, Paterna, Valencia, Spain; Translational Bioinformatics Unit, Navarrabiomed, Complejo Hospitalario de Navarra (CHN), Universidad Pública de Navarra (UPNA), IdiSNA, Pamplona, Spain; Technische Universität Dresden, Institute of Waste Management and Circular Economy, Pirna, Germany; Bioclear Earth B.V., Groningen, The Netherlands; Brandenburgische Technische Universität Cottbus-Senftenberg, Chair of Circular Economy, Cottbus, Germany; Institute for Integrative Systems Biology I2SysBio (University of Valencia - CSIC), Paterna, Spain

**Keywords:** Metagenomics, anaerobic digestion, biogas, SAOB, MBA03, *Darwinibacteriales*

## Abstract

Biogas, a mix of CO_2_, CH_4_ and small proportions of other gases, is a biofuel obtained by anaerobic digestion (AD). Biogas production is often considered a black box process, as the role and dynamics of some of the microorganisms involved remain undisclosed. Previous metataxonomic studies in the frame of the MICRO4BIOGAS project (www.micro4biogas.eu) revealed that MBA03, an uncharacterised and uncultured bacterial taxon, was very prevalent and abundant in industrial full-scale AD plants. Surprisingly, no culturable specimen or genome of this taxon has ever been reported, so its role in AD has remained unclear. In the present work, thirty samples derived from anaerobic digesters were sequenced, allowing the reconstruction of 108 metagenome-assembled genomes (MAGs) potentially belonging to MBA03. According to phylogenetic analyses and genomic similarity indices, MBA03 constitutes a new bacterial order, proposed as *Darwinibacteriales* ord. nov., which includes *Darwinibacter acetoxidans* gen. nov., sp. nov. of the family *Darwinibacteriaceae* fam. nov., along with *Wallacebacter cryptica* gen. nov., sp. nov. of the *Wallacebacteriaceae* fam. nov. Ecotaxonomic studies determined that AD processes are the main ecological niche of *Darwinibacteriales*. Moreover, metabolic predictions identified *Darwinibacteraceae* members as putative syntrophic acetate oxidising bacteria (SAOB), as they encode for the reversed Wood-Ljungdahl (W-L) pathway coupled to the glycine cleavage system. This suggests that *Darwinibacteraceae* members work in collaboration with hydrogenotrophic archaea to produce methane in industrial biogas plants. Overall, our findings present *Darwinibacteriales* as a potential key player in anaerobic digestion and pave the way towards the complete characterisation of this newly described bacterial taxa.

## 1 Introduction

The evidence supporting the need for a transition from fossil to renewable energy is overwhelming and unanimous (Yaqoob et al., 2021). In recent years, biogas has arisen as one of the leading alternatives to non-renewable fossil fuels such as natural gas. Biogas is produced by anaerobic digestion (AD) of organic material, through a complex microbiological process in which several groups of microorganisms with diverse metabolic abilities work in close collaboration to obtain methane as the main product (Schnürer, 2016). In addition, biogas production contributes to the utilisation of today’s global waste streams of various origins as a source of organic matter (Tsang et al., 2019).

Biogas primarily consists of methane (CH_4_), carbon dioxide (CO_2_) and other gases like sulfuric acid (H_2_S), nitrogen (N_2_), ammonia (NH_3_), and hydrogen (H_2_) in lower concentrations (Yaqoob et al., 2021). Despite the complexity of the AD process, the microbial metabolic reactions that occur in this process can be summarized in four key steps: hydrolysis, acidogenesis, acetogenesis and methanogenesis (Schnürer, 2016). Briefly, during hydrolysis, complex macromolecules are broken down into various oligo-, di- and monomers, the final concentrations of which will depend on the biomass source (Nwokolo et al., 2020). These molecules are uptaken and used by acidogenic bacteria to produce H_2_, CO_2_, alcohols and volatile fatty acids (VFAs) during acidogenesis, while in acetogenesis, acetogenic bacteria synthesize acetic acid through the Wood-Ljungdahl (W-L) pathway or autotrophically from H_2_ and CO_2_. Subsequently, methanogens metabolize acetate, hydrogen and/or methylated compounds to produce methane, creating an equilibrium and a co-dependence between bacteria and archaea (Vanwonterghem et al., 2016; Nwokolo et al., 2020). Two pathways can occur in the production of methane from acetate: the first one, directly by acetoclastic methanogens, which split acetate to methane and CO_2_, with the methyl group yielding methane and the carboxyl group yielding CO_2_. The other pathway results from syntrophic acetate-oxidizing bacteria (SAOB) oxidizing acetate to CO_2_ and H_2_, which is subsequently used by hydrogenotrophic archaea to obtain methane. Thus, this pathway must be coupled with hydrogenotrophic methanogenesis (SAO-HM).

Despite the high economic potential, knowledge for the production of biogas with AD has remained stagnant, as the high variability of both the different operational parameters and the native microbial communities involved represent a challenge for its optimization and standardization. Improving the feasibility of the process and achieving a consistent and controlled way to obtain biogas would constitute a huge change in the energy industry (Nielsen & Angelidaki, 2008). But reaching such an ambitious goal requires gaining insights in the microbial machinery behind AD and methanogenesis, and this goes through the study of a variety of AD systems with different chemical, taxonomical and operational parameters.

In this context, the EU-funded project MICRO4BIOGAS (https://micro4biogas.eu) was born in 2021. The ultimate objective of the project is to improve the yield, speed, quality and robustness of biogas production by optimising the AD microbiome. In the frame of the project, 80 different samples were collected from various full-scale AD plants in the Netherlands, Germany and Austria.

After metataxonomic analyses of the 80 samples during MICRO4BIOGAS project, a miscellaneous taxonomic group (i.e., MBA03) was found to be very prevalent and abundant, with some samples reaching abundances up to 38% of total bacteria. Moreover, MBA03 was among the taxa that correlated most strongly with operational (i.e., temperature) and chemical parameters (i.e., acetic acid) (Otto et al., 2023), suggesting that it may be a microorganism of great relevance for the AD process. MBA03 (AB114313.1) was described for the first time by Tang et al. as an uncultured clone obtained from a thermophilic anaerobic municipal solid-waste digester (Tang et al., 2004). The proposal of MBA03 exclusively relied on the 16S rRNA gene sequence. Since then, a series of metataxonomic studies have reported the presence of MBA03 in anaerobic digester systems (Dyksma et al., 2020; FitzGerald, 2018; Laguillaumie et al., 2022), representing up to 70% of the relative abundance in some samples (FitzGerald, 2018). This taxon has been associated with a higher production of methane and has been suggested as an indicator of the stability of the AD process (Fang et al., 2022; Laguillaumie et al., 2022).

Surprisingly, MBA03 has not been properly taxonomically delimited and no culturable specimen or genome of this taxon has ever been reported. However, according to SILVA database (SSU 138.1 Ref NR), MBA03 represents a heterogenous group of more than 150 16S rRNA sequences. Therefore, a deeper characterization of this taxonomic group was urgently needed for unravelling its role in AD processes.

In this work, we used metagenomic data from 30 MICRO4BIOGAS samples and other AD studies to propose MBA03 as a member *Darwinibacteriales* ord. nov., a not-yet-cultured bacterial taxa belonging to the class *Clostridia. Darwinibacteriales* includes two different genera: *Darwinibacter acetoxidans* gen. nov., sp. nov., of the *Darwinibacteriaceae* family, and *Wallacebacter cryptica* gen. nov., sp. nov., of the *Wallacebacteriacea* fam. nov. Based on our results, both *Darwinibacter* and *Wallacebacter* predominantly inhabit AD environments. Furthermore, we have inferred that *Darwinibacter* may play a key role in AD processes related to biogas production, as it includes the reversed Wood-Ljungdahl pathway coupled to the glycine cleavage system, which is required for the synthesis of H_2_ and CO_2_ from acetate and the subsequent production of methane by hydrogenotrophic methanogenesis.

## 2. Materials and methods

### 2.1. Sample description, DNA extraction and metagenomic sequencing

As part of the MICRO4BIOGAS project, a total of 80 samples were collected from anaerobic digestion systems at 45 large-scale reactors across Germany, Austria and the Netherlands (Otto et al., 2023). A subset of 30 samples was analysed through metagenomic sequencing. This subset included samples enriched in MBA03, hydrolytic, acidogenic and acetogenic bacteria and/or methanogenic archaea. A full description of the samples can be found in Supplementary Table 1.

The sludge samples were washed with PBS, and the DNeasy® PowerSoil® Pro kit (QIAGEN, Germany) was used for DNA extraction. The metagenomic DNA was fragmented, end polished, A-tailed, and ligated with Illumina adapters. Libraries were sequenced at Novogene (UK) using the NovaSeq 6000 Illumina platform (150 bp x 2). The average sequencing depth was 45M reads/sample (min.: 35M reads; max.: 55M reads). Additional information about DNA extraction and sequencing can be found at Supplementary Methods.

### 2.3. Metagenomic analysis

Reads were filtered and quality checked as described at Supplementary Methods. MEGAHIT (v1.2.9) was used with default parameters for assembling the filtered reads (D. Li et al., 2015). Two different binning strategies were applied: MetaBAT (Kang et al., 2019) and MaxBin (Wu et al., 2016). The MAGs obtained were refined with Das Tool (Sieber et al., 2018) and classified according to their quality with CheckM (Parks et al., 2014): high quality (HQ, completeness ≥ 90% and contamination ≤ 5%), good quality (GQ, completeness ≥ 80% and contamination ≤ 10%) and medium-low quality (LQ, completeness ≤ 80% and/or contamination ≥ 10%), following the criteria of Bowers et al., 2017 and Feng et al., 2021. After that, Centrifuge (Kim et al., 2016) was used to study the taxonomic profile of the samples, while the resulting MAGs were taxonomically annotated with Microbial Genome Atlas (MiGA) (Rodriguez-R et al., 2018).

### 2.4. Recovery of MBA03 genomes via phylogenetic analyses

First of all, 16S rRNA sequences were extracted from the high-quality MAGs using Prokka (v1.14.6) (Seemann, 2014). These sequences were aligned together with the 157 available MBA03 16S rRNA sequences from SILVA database (SSU 138.1 Ref NR) with the SINA Aligner (v1.2.11) (Pruesse et al., 2012), and afterwards a phylogenetic tree was computed with FastTree (v2.1.1) (Price et al., 2009) and visualized with the Interactive Tree Of Life (iTOL) (v5, Letunic & Bork, 2021). Only the MAGs whose sequences formed a monophyletic group with MBA03 were selected for further analysis. Due to the limitations associated to the binning process, not all MAGs contained 16S rRNA sequences, while others contained more than one sequence, so not all the MAGs corresponding to MBA03 could be identified by 16S rRNA.

### 2.5. MBA03 identification from phylogenomic data

After identifying the first MAGs hypothetically belonging to MBA03, a two-step approach was followed to recover the remaining MAGs belonging to the MBA03 taxonomic group which did not contain rRNA operons.

As a first step, all the MAGs whose 16S rRNA gene formed a monophyletic group with MBA03 according to the phylogenetic tree were annotated using the whole genome classifier tool Genome Taxonomy Database toolkit (GTDB-tk; v2.1.1) (Chaumeil et al., 2022). Although GTDB does not contain any genome identified as “MBA03”, the resulting annotation was used as a first hypothesis to elucidate which of the genomes actually corresponded to MBA03.

As a second step, all the HQ and GQ MAGs recovered in the project (n = 1053) were also annotated GTDB-tk. In addition, a phylogenomic tree was constructed using the UBCG pipeline (v3.0) (Na et al., 2018). Once all the MAGs corresponding to MBA03 were identified, genomes belonging to the MBA03 order in the GTDB were downloaded and included in the dataset. Afterwards, genomes from the Biogasmicrobiome repository (https://biogasmicrobiome.env.dtu.dk/), and from (Dyksma et al., 2020) were annotated and incorporated into the dataset.

### 2.6. Taxonomic delimitation of the MBA03 taxonomic clade

Prokka (v1.14.6) (Seemann, 2014) was used again to obtain all the 16S rRNA sequences from the MAGs and genomes identified as potential MBA03 representatives according to the previous phylogenomic analyses. To better study the phylogenetic patterns, 16S rRNA sequences corresponding to the most closely related taxonomic classes were set as outgroups. Sequences from class *Clostridia* and *Limnochordia* were downloaded from the Living Tree Project (LTP) and dereplicated using cd-hit (4.8.1) (W. Li & Godzik, 2006) at a 0.97 similarity threshold. Sequences were aligned with SINA Aligner (v1.2.11) (Pruesse et al., 2012) and incorporated into the previous dataset. Finally, a phylogenetic tree was computed with FastTree (Price et al., 2009), and visualized with iTOL (v5; Letunic & Bork, 2021). The reference sequence of the bacterial clone identified for the first time as MBA03 was also downloaded from the NCBI (AB114313.1) and included in the tree.

Following the same procedure as for the phylogenetic tree, genomes from *Clostridia* and *Limnochordia* were downloaded from RefSeq NCBI Assembly Database, filtering by “Assembly from Type Material” and “Complete Genomes” (Supplementary Table 2). These genomes were analysed altogether with the genomes identified as MBA03 and a phylogenomic tree was created with UBCG (Na et al., 2018) and visualized with iTOL (v5; Letunic & Bork, 2021).

### 2.7. Analysis of the ecological niche of MBA03

In order to explore the ecological niches of MBA03, an analysis with IMNGS (v1) (Lagkouvardos et al., 2016) was carried out using a 97% identity cut-off. The same sequences used for the phylogenetic analysis were dereplicated using cd-hit (v4.8.1) (W. Li & Godzik, 2006) at a 0.9 identity threshold. IMNGS results were filtered with a series of in-house Python scripts and manual curation (https://github.com/roserpr/IMNGSanalysis.git). More information about the filtering process can be found at Supplementary Methods.

In order to determine the abundance of MBA03 among the samples, coverM (v0.5.0, https://github.com/wwood/CoverM) was used, comparing the raw reads of the 30 sequenced reactors against the genomes assigned to MBA03.

### 2.8. Functional annotation

MAGs belonging to MBA03 with the higher qualities were selected (completeness > 90% and contamination limited to < 3%, both calculated with CheckM (Parks et al., 2014)). These MAGs were annotated with Bakta (Schwengers et al., 2021) and they were analysed with Kofamscan (Aramaki et al., 2020) and KEGG-Decoder (Graham et al., 2018).

As mentioned above, acetate as an intermediate is of particular importance during anaerobic reactions, especially for SAOB bacteria and hydrogenotrophic methanogenesis. In order to elucidate if MBA03 was a SAOB, the protein sequences of all the enzymes involved in the reversed WL pathway and the reversed WL pathway coupled to the glycine cleavage system (GCS) (Supplementary Table 3) were downloaded from UniProt (Coudert et al., 2023) (Swissprot), narrowing to all bacteria (taxonomy_id:2), and a database was created with Diamond (Buchfink et al., 2015). A Diamond BLASTp of the annotated protein sequences was performed against this database to detect presence and absence of each gene in all MAGs with 50% coverage and identity thresholds. For the whole metabolic analysis, any enzyme found in more than 70% of the MAGs of each family was considered to be present in the genome.

## 3. Results and discussion

### 3.1. Genomic reconstruction

MBA03 has proven abundant in a variety of anaerobic digesters (Dyksma et al., 2020; Liu et al., 2023). Given its importance in AD environments, this work focused on the recovery and characterisation of the genome of this taxonomic group using 30 metagenomic datasets derived from the MICRO4BIOGAS project and other genomic data available in public databases.

After metagenomic sequencing, assembly and binning of sludge samples, the MAGs obtained were classified according to their quality, and 1053 MAGs (HQ and GQ) were chosen for further analysis. After the annotation of the MAGs with MIGA and the metagenomes with Centrifuge, in neither case MBA03 was reported despite being a highly abundant taxon according to the metataxonomic results (Otto et al., 2023). This was expected as genomic databases did not include MBA03 genomes, probably reflecting a lack of consensus on the nomenclature of this taxonomic group.

### 3.2. Taxonomic analysis of MAGs

In order to identify the genomes belonging to MBA03, a phylogenetic tree was constructed using the 16S rRNA sequences extracted from the HQ MAGs and 157 sequences classified as MBA03 according to the SILVA database (SSU 138.1 Ref NR) (Supplementary Figure 1). The sequences from SILVA formed a monophyletic clade, and only 16 sequences from the metagenomic dataset fell within this subtree. The MAGs corresponding to these 16 sequences were annotated with GTDB-tk (v2.1.1) and 12 out of the 16 MAGs shared the same annotation: they belonged to phylum Bacillota, class *Limnochordia*, order DTU010. From here, two different families were identified: DTU010 and DTU012 (Supplementary Table 4).

After this preliminary analysis, the HQ and GQ MAGs (n=1053) were annotated with GTDB-tk (Supplementary Table 5). A total of 45 MAGs were classified as members of the order DTU010, and they were subsequently treated as potential representatives of MBA03. The phylogenomic tree of HQ and GQ MAGs (Figure 1) also revealed a large proportion of MAGs belonging to *Bacillota* (n = 503), while 104 MAGs were classified as *Bacteroidota,* 77 as *Actinobacteriota,* 63 as *Verrucomicrobiota* and 32 as *Chloroflexota,* while 80 MAGs corresponded to Archaea. These results were expected considering previous studies reporting the average AD microbiome (Sundberg et al., 2013; Abendroth et al., 2015; Treu et al., 2016; Kirkegaard et al., 2017; Otto et al., 2023). Moreover, it was confirmed that MBA03 formed a monophyletic clade with all 45 sequences annotated as DTU010. Two different subclades could be clearly distinguished, corresponding to families DTU010 and DTU012 according to the GTDB taxonomy.

**Figure 1.**
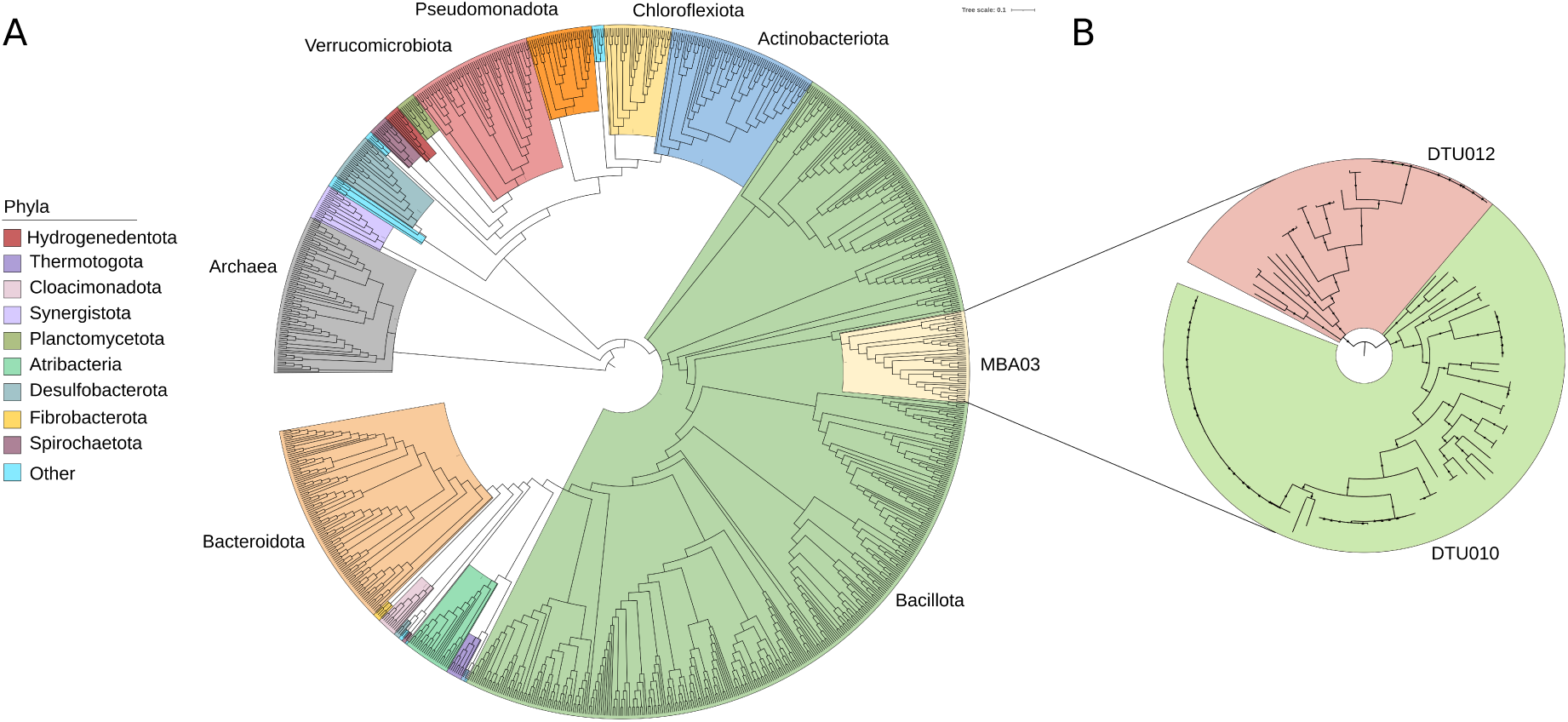
(A) Dendrogram of all the HQ and GQ MAGs from MICRO4BIOGAS (n=1053), computed with UBCG and visualized with iTOL (v5). The legend shows a classification by different colours corresponding to different phyla, according to the annotation obtained with GTDB-tk (v2.1.1), and a highlight on MBA03 (within *Bacillota*). (B) Tree with 108 leaves, computed with FastTree and visualized with iTOL (v5), corresponding to the MAGs belonging to order DTU010 according to GTDB-tk. Two clades corresponding to two families (DTU010 in orange and DTU010 in yellow) can be seen, for which the names *Darwinibacteriaceae* and *Wallacebacteriaceae* are proposed, respectively. In both figures, bootstrap values greater than 0.75 are represented by a small triangle in each branch.

For the enlargement of the dataset, 2237 MAGs from the Biogasmicrobiome repository (https://biogasmicrobiome.env.dtu.dk/), 16 from Dyksma et al. and 19 from GTDB (release 207) were downloaded and annotated with GTDB-tk (Dyksma et al., 2016). After filtering, a total of 63 potential MBA03 MAGs were compiled. Together with the previous 45 MAGs (HQ and GQ) retrieved from MICRO4BIOGAS data, the final dataset was composed of 108 genomes. These MAGs were used for creating a phylogenomic tree (Figure 2), leading to an initial taxonomic delimitation of MBA03: (i) there were two families, named DTU010 (which accounted for 82 MAGs) and DTU012 (with 26 MAGs) according to GTDB database; (ii) DTU010 clade included 20 different taxa annotated in GTDB-tk. The most common was DTU010 sp. 002391385, followed by sp. 900018335, and both formed monophyletic clades; and (iii) there were 10 different taxa belonging to DTU012, with the most common species being DTU012 sp.900019385.

**Figure 2.**
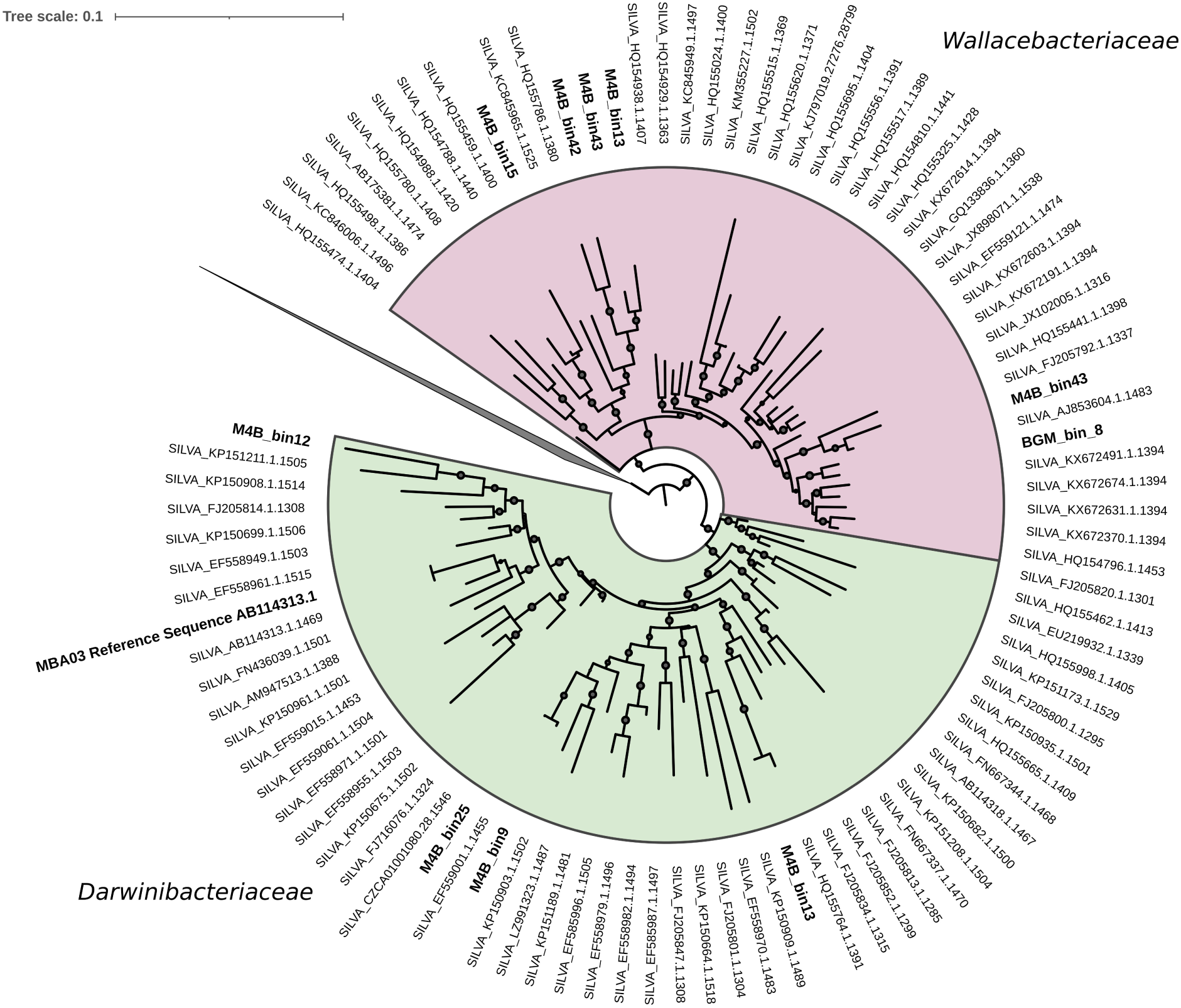
Phylogenetic tree based on 16S rRNA sequence from our own database and from SILVA SSU 138.1 Ref NR database. The sequences in bold letter belong to the MAGs from our database, while the other sequences are the ones downloaded from SILVA. Output groups were collapsed for a better visualization of the MBA03 clade.

Thereby, it was proposed the new order to be named *Darwinibacteriales* ord. nov., and the two families *Darwinibacteriaceae* fam. nov., and *Wallacebacteriaceae* fam. nov., after Charles Darwin and Alfred Wallace, who are considered the fathers of the Theory of Evolution by natural selection.

### 3.3. Final taxonomic delimitation of MBA03

According to these results, the order *Darwinibacteriales* ord. nov. (DTU010 according to GTDB) was hypothesized to be equivalent to MBA03 (SILVA’s name), but further phylogenetic evidence was needed to get a proper delimitation of the clade. For that reason, phylogenetic analyses based on the 16S rRNA gene were carried out. A total of 52 16S rRNA sequences were identified by Prokka (Seemann, 2014) in the 108 MAGs (i.e., not all MAGs contained 16S rRNA sequences), and they were computed together with the 157 16S rRNA sequences downloaded from the SILVA (SSU 138.1 ref NR) database in a phylogenetic tree (n=209). Moreover, 772 sequences from *Clostridia* were added as outgroups.

The phylogenetic tree (Figure 2) confirmed the abovementioned hypothesis: the sequences annotated as MBA03 in SILVA formed a monophyletic clade with a clear subdivision of two clades, which included two distinct families (*Darwinibacteraceae*/DTU010 and *Wallacebacteriaceae*/DTU012). Clade I was formed by 52 sequences and included the MBA03 reference sequence, whereas Clade II accounted for 44 sequences.

It is worth highlighting that, although both *Hydrogenispora* and *Darwinibacteriales* (MBA03) are annotated as class *Limnochordia* in the SILVA database, *Hydrogenispora* is annotated as *Clostridia* in the LPSN (List of Prokaryotic names with Standing in Nomenclature, Parte et al., 2020), which provides the internationally accepted and regulated taxonomic classification. This annotation, supported by other authors (FitzGerald, 2018; Westerholm et al., 2016), was confirmed once the tree was computed, as both *Hydrogenispora* and *Darwinibacteriales* appeared to be two different orders from class *Clostridia*. In other words; *Darwinibacterales* is proposed as a new order that belongs to class *Clostridia*.

Finally, a phylogenomic tree (Figure 3) was computed with the 108 genomes classified as MBA03 together with 206 genomes from *Clostridia* and one genome from *Limnochordia* as outgroups (Supplementary table 2). This confirmed the hypothesis that MBA03 belongs to a new order within class *Clostridia,* and it comprises two families.

**Figure 3.**
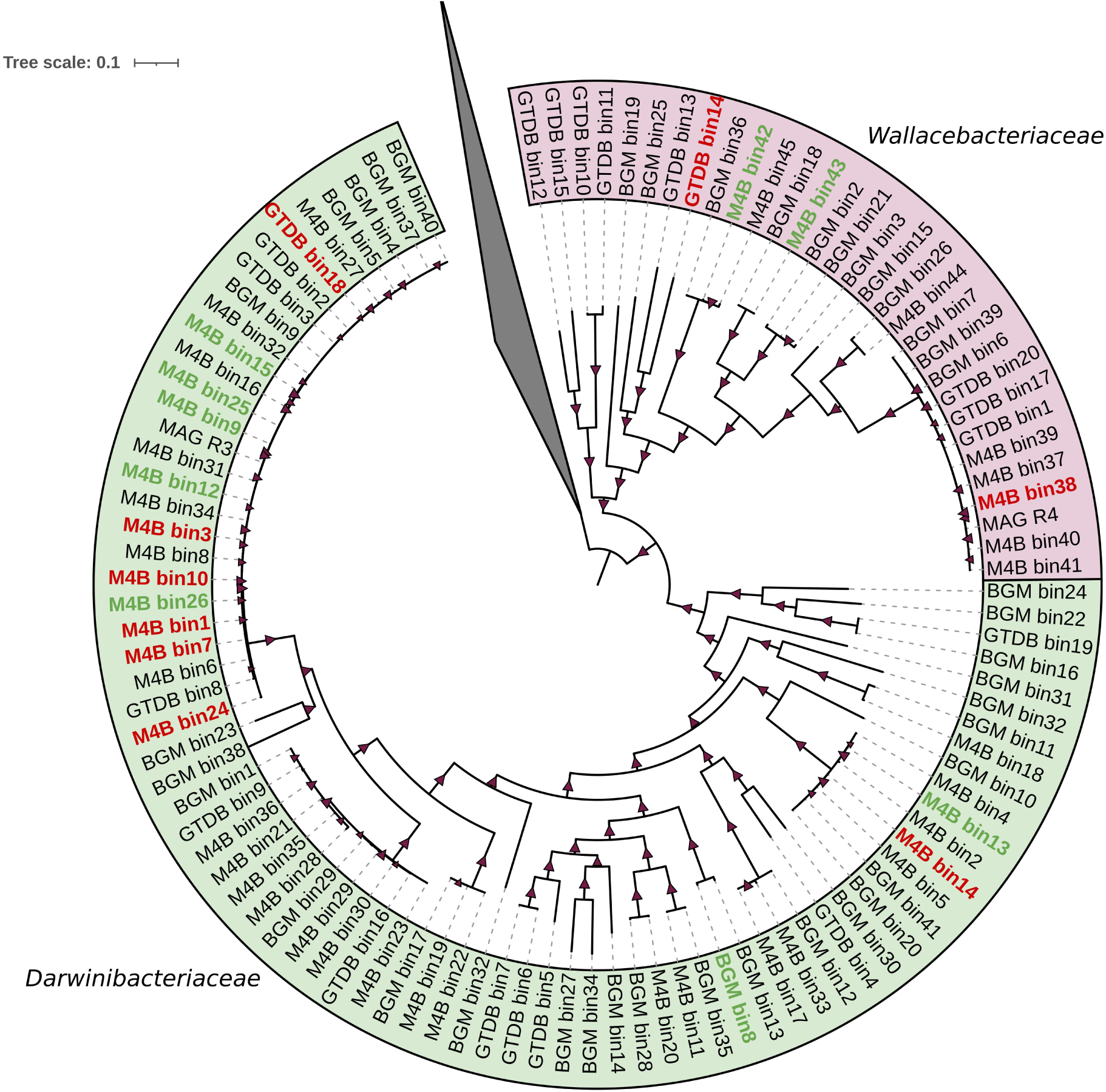
Phylogenomic tree showing only the *Darwinibacteriales* (MBA03 in SILVA) clade, which includes two potential new families: *Darwinibacteriaceae* (DTU010) shaded in pink and *Wallacebacteriaceae* (DTU012) shaded in green. MAGs in green (n=9) belong to *Darwinibacteriales* (MBA03), as there is evidence at the genetic and genomic level. MAGs in red (n=9) may be MBA03, but their 16S rRNA phylogeny is not concordant. This could be explained by errors during the binning process. The remaining MAGs (n=90) are incomplete and/or 16S rRNA sequences could not recovered from the MAGs. Although they form a cluster with the rest of the MBA03 MAGs, they should remain as hypothetical members of this order. Output groups were collapsed for a better visualization of the MBA03 clade.

According to these results, there is sufficient evidence to confirm that MBA03 belongs to a new taxonomical order, *Darwinibacteriales* ord. nov., which includes two potential new families, *Darwinibacteriaceae* fam. nov., (DTU010 according to GTDB) and *Wallacebacteriaceae* fam. nov.,(former DTU012). It is important to note that samples from MICRO4BIOGAS provided the largest number of MAGs belonging to MBA03, meaning that the samples and datasets from this project marked a significant breakthrough towards the genetic isolation of MBA03 genomes. The formal description of these taxa can be found in Supplementary Material.

Once the two families from the order *Darwinibacteriales* (*Darwinibacteriaceae* and *Wallacebacteriaceae*) were properly delimited, the read coverage of both families in the samples was calculated. 10 MAGs from DTU010 and 4 MAGs from DTU012 were used as reference for read mapping. The results showed that DTU010, with a mean abundance of 4.1% and a maximum abundance of near 18%, was considerably more abundant than DTU012, with a mean abundance of 0.6% (Supplementary Table 6).

### 3.5. Ecological distribution *Darwinibacteriales*

The ecological distribution of *Darwinibacteriales* was evaluated using IMNGS. This tool quantifies the prevalence and abundance of any 16S rRNA sequence of interest introduced as input, giving information about the number of coincidences with the samples included in its database. IMNGS also informs about the type of sample (e.g., soil, water, anaerobic digester, environmental) showing a hit with the input sequence. A total of 8856 SRA accessions containing 16S rRNA sequences from *Darwinibacteriales* were detected according to sequence similarity. Due to the high amount of data obtained, only the samples with an abundance of *Darwinibacteriales* higher than 0.1% were considered at first (Figure 4A). An exhaustive analysis of the sample metadata revealed that more than 99% of them belonged to anaerobic digestion studies, meaning that *Darwinibacteriales* is a very specialized bacterial group that grows and develops in this specific environment. This result reinforces the hypothesis that MBA03 *Darwinibacteriales* may play a central role in the AD process.

**Figure 4.**
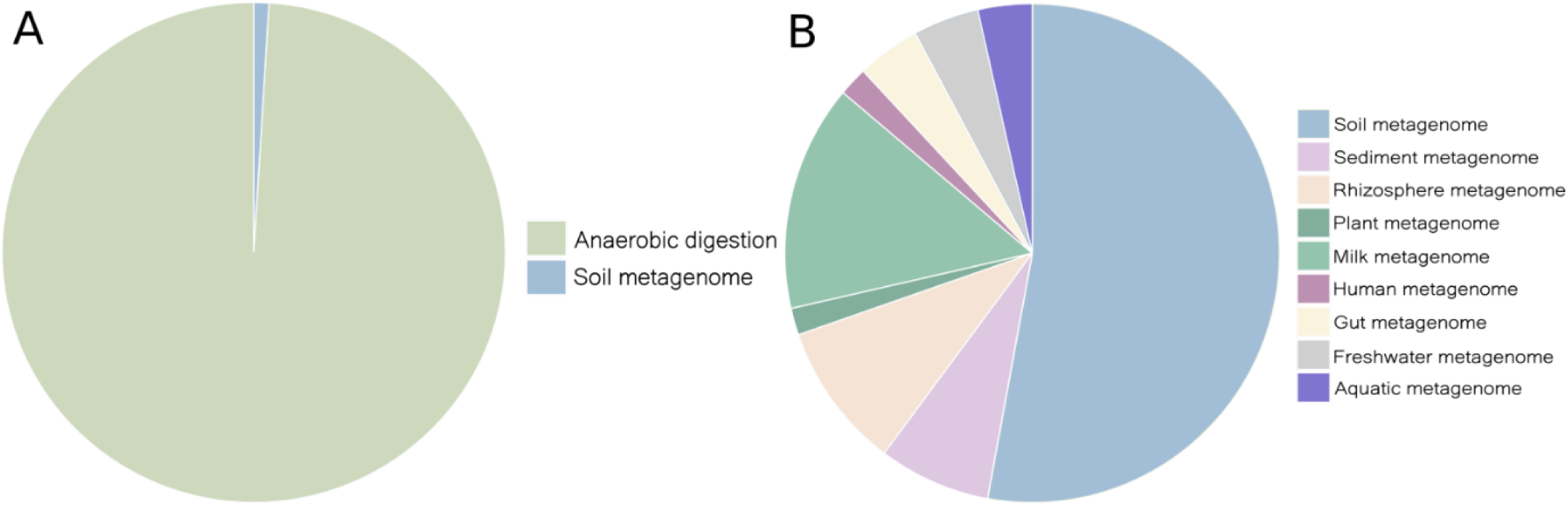
(A) Ecological niches of *Darwinibacteriales* based on coincidences of 16S rRNA sequences with samples of the IMNGS database with an abundance above 0.1%, (B) Ecological niches of *Darwinibacteriales* according to the samples with an abundance below 0.1% and excluding all the samples related to anaerobic digestion for a better insight into the wide range of environments from where it may be introduced to anaerobic reactors.

To study the ecological niches besides anaerobic digestion, a similar analysis was performed considering only the samples with an abundance of *Darwinibacteriales*-related sequences below 0.1%. The resulting 8650 hits were then filtered in two steps: (i) first, those related to anaerobic digestion were discarded, and (ii) results with less than 5 hits in total were removed from the dataset, leaving 1636 samples that were categorised into different groups (Figure 4B).

Soil was the second most common environment in which MBA03 could be found, followed by rhizosphere and sediment samples. However, the abundance of *Darwinibacteriales* in these habitats was substantially lower than in the case of anaerobic digesters. Therefore, it can be hypothesized that these bacteria are found in soil, plants and grounds, in a quiescent or inactive form. When agricultural residues, soils or sludges are taken as substrate for anaerobic digestion, bacteria from the *Darwinibacteriales* order thrive in the AD environment. Similarly, the presence of MBA03 in gut and milk microbiomes can be explained by the dietary habits of the animals. As *Darwinibacteriales* exists in the soil, herbivorous and omnivorous animals may ingest this group of bacteria during their feeding. Another hypothesis would be that *Darwinibacteriales* is part of the native gut microbiota of the animals, but at a very low abundance, and would thus be a reservoir which would guarantee the prevalence in AD plants fed with those substrates. Therefore, when manure is used as a substrate, *Darwinibacteriales* would be introduced into the bioreactors.

### 3.6. Unveiling the functional potential of *Darwinibacteriales*

In anaerobic digesters, the metabolic diversity of *Darwinibacteriales* remains challenging due to the absence of known cultivable representatives. A total of 45 and 16 MAGs from *Darwinibacteriaceae* (former DTU010) and 16 from *Wallacebacteriaceae* (former DTU012), respectively, were analysed to gain insights into the potential metabolic pathways of this bacterial order (Supplementary Table 7).

*Darwinibacteriales* are chemoorganoheterotrophs that derive energy from anaerobic respiration through oxidative phosphorylation. They utilize inorganic molecules of ferric iron (Fe3+) as potential electron acceptors (K02012 and K02011). These bacteria also possess sulphide:quinone oxidoreductase (EC:1.8.5.4), an enzyme involved in the oxidation of sulphides, including hydrogen sulphide, enabling them to be chemolitotrophic. Their ability to grow on H_2_S is interesting, as H_2_S is usually not oxidized under anaerobic conditions. However, there are some exceptions.

*Darwinibacteriaceae* and *Wallacebacteriaceae* have complete sets of genes for glycolysis (Embden-Meyerhof pathway; M00001), gluconeogenesis (M00003), and pyruvate oxidation (M00307) pathways. However, they lack certain essential genes required for the tricarboxylic acid (TCA) cycle, specifically isocitrate dehydrogenase. Both bacterial families can convert glycogen to glucose-6P.

MAGs from both families contain genes encoding beta-N-acetylhexosaminidase for bacterial cell wall peptidoglycan degradation and recycling, pullulanase for starch degradation, and beta-glucosidase for cellobiose metabolism. *Darwinibacteriaceae* members also possess additional genes encoding chitinase for chitin degradation, D-galacturonate isomerase for pectin utilization, and alpha-amylase for starch hydrolysis. *Wallacebacteriaceae* members carry the gene encoding oligogalacturonide lyase, allowing them to degrade pectin. Both *Wallacebacteriaceae* and *Darwinibacteriaceae* exhibit the ability to degrade catechol and methylcatechol through catechol 2,3-dioxygenase enzymes (EC 1.13.11.2). However, the presence of these compounds can be toxic to microorganisms in anaerobic digesters, inhibiting the digestion process. These findings suggest that each bacterial family can exploit different organic substrates as carbon and energy sources, maybe facilitating their coexistence. This also indicates that the importance of MB03 does not only restrict to one single phase of anaerobic digestion. Apart from acetogenesis and syntrophic acetate oxidation, MB03 could also be involved in the hydrolysis phase as well.

Members of *Darwinibacteriales* assimilate nitrogen by converting NH_3_ to organic nitrogen through the enzymatic activities of glutamine synthetase (EC 6.3.1.2) and glutamate (NADPH) synthase (EC 1.4.1.13). However, genes associated with other nitrogen metabolism processes, such as nitrate reduction, denitrification, nitrogen fixation, nitrification, and anaerobic ammoniumoxidation, were not observed. Most representatives of *Darwinibacteriaceae* have auxotrophic requirements for various amino acids, while *Wallacebacteriaceae* representatives are auxotrophic for a smaller set of amino acids (Supplementary Table 7).

Both bacterial families share several ABC transporters, including ABCB-BACATP binding cassettes and NitT/TauT family transporters. They also possess multiple sugar transport systems. *Darwinibacteriaceae* exhibits ABC transporters for iron (III), phosphonate, alpha-glucoside, aldouronate, sn-glycerol 3-phosphate, cobalamin, and haemolysin transport systems. Both *Darwinibacteriaceae* and *Wallacebacteriaceae* show transport systems for maltose/maltodextrin, arabinogalactan/maltooligosaccharide, raffinose/stachyose/melibiose, glucose/mannose, trehalose/maltose, cellobiose and other tansport systems (See Supplementary Table 7).

Members of *Darwinibacteriales* possess resistance genes against vancomycin (vanW) and tetracycline (tetM and tetO). They also encode multidrug efflux pumps (mdlA/smdA; mdlB/smdB; efrA/efrE) to withstand various antimicrobial compounds.

Both families are motile, as indicated by the presence of flagella synthesis genes. However, *Darwinibacteriaceae* lacks genes for the MS/C ring type III secretion system. Members of both taxa possess the enzymatic machinery for chemotactic movement and detection and transport of D-ribose and D-galactose via ABC transporters.

### 3.7 Members of *Darwinibacteriaceae* as potential SAOB

Members of the MBA03 lineage were previously hypothesised to be SAOB in AD based on correlation studies (Zheng et al., 2019; Zeng et al., 2021; Perman et al., 2022); however, the metabolism of this bacterial lineage, classified in this study as *Darwinibacteriaceae*, remains poorly understood and characterised due to the lack of cultured representatives. Recently, the co-occurrence of members of *MBA03* with other potential SAO bacteria, such as DTU014 and *Syntrophaceticus*, and with hydrogenotrophic methanogens such as *Methanothermobacter*, *Methanoculleus* and *Methanosarcina,* has been observed (Otto et al., 2023). This suggests that several taxonomic groups identified as potential SAOBs, including *Darwinibacteriaceae*, grow simultaneously with methanogens to provide a continuous supply of H_2_ and CO_2_ for hydrogenotrophic methanogenesis. Usually, hydrogenotrophic methanogens are associated with high performance plants, whereas the acetoclastic methanogens *Methanothrix* (formally *Methanosaeta*) are found specifically in low performance systems such as sewage sludge digesters (Sunderberg et al. 2013; Abendroth et al., 2015). Therefore, due to the co-occurrence between members of *Darwinibacteriaceae* and hydrogenotrophic methanogens, we hypothesise that the presence of *Darwinibacteriaceae* could be associated with a higher biogas production in industrial AD plants, although further analyses are required.

It has been seen that SAO bacteria possess either the known reversed WL pathway or an alternative route coupled to the GCS (Zeng et al., 2021). To confirm whether members of the *Darwinibacteriaceae* and *Wallacebacteriaceae* are SAOBs, their MAGs were mined to identify genes of the reversed WL pathway and the GCS (Figure 5). In all members of the *Darwinibacteriaceae* family, gene *Acka* was consistently detected, whereas *Pta*, *Acs*, the *CODH* complex, and *MetF* were conspicuously absent. Most of the *Darwinibacteriaceae* MAGs contained *FolD*, *Fhs*, and *Fdh* enzymes, which play vital roles in the acquisition of CO_2_ and H_2_ from formate. The absence of gene *Pta* and *Acs* could be justified by the presence of glycine reductase (*Grd*) in the glycine cleavage system, which facilitates the conversion of glycine for entry into the GCS (Li et al., 2022). Thus, the entire reverse WL-GCS pathway was encoded by members of *Darwinibacteriaceae*, with only two genes, namely *Pta* and *Acs* missing, which are in fact not necessary for the completion of the pathway, as in the GCS, CH3-CO-Pi is directly converted to glycine through the function of glycine reductase (Grd) (Li et al., 2022). The remaining enzymes were identified in more than 70% of the *Darwinibacteriaceae* MAGs, with the exception of *Dld*, which was present in only 60% of them.

**Figure 5.**
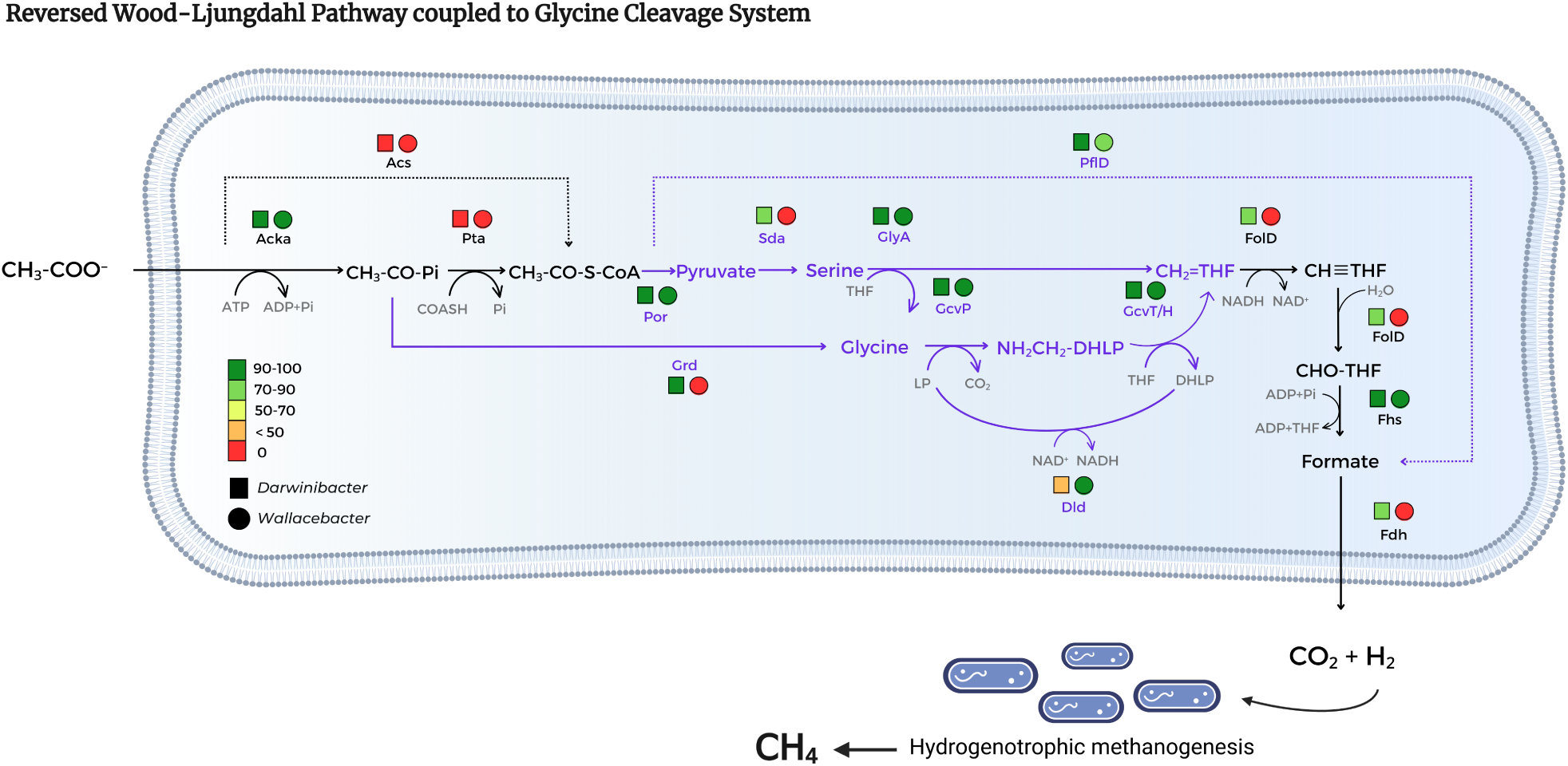
Mining of reverse WL pathway (black lines) coupled with glycine cleavage system (purple lines) (WL-GCS) genes in MAGs of the families *Darwinibacteriaceae* (n=45) and *Wallacebacteriaceae* (n=16). Red to green colors indicate the percentage of MAGs in which the genes were identified. A gene was considered to be present in each family if it was found in more than 70% of the MAGs. Bacteria from the family *Darwinibacteriaceae* encode the genetic repertoire necessary to metabolise acetate to produce CO2 and H2, which are then used by hydrogenotrophic archaea to produce methane, whereas *Wallacebacteriaceae* showed a truncated pathway. Abbreviations: *AckA*; acetate kinase, *Pta*; Phosphate acetyltransferase, *Acs*; Acetyl-coA synthetase, *Por*; pyruvate-ferredoxin reductase, *Sda*; L-serine dehydratase, *GlyA*; glycine hydroxymethyltransferase, Grd; glycine reductase, *GcvP*; glycine dehydrogenase, *GcvH/T*; glycine cleavage system H/T protein, *Dld*; dihydrolipoyl dehydrogenase, *PflD*; pyruvate-formate lyase, *AcsE;* iron sulfur protein methyltransferase, *metF*; methylenetetrahydrofolate reductase, *FolD*; methylenetetrahydrofolate dehydrogenase/cyclohydrolase, *Fhs*; formate-tetrahydrofolate ligase, *Fdh*; formate dehydrogenase.

In the case of the family *Wallacebacteriaceae*, only genes *AckA* and *Fhs* were identified, while the other genes of the pathway were absent. The absence of critical enzymes of the reversed Wood-Ljungdahl pathway, including *Sda*, *Grd*, *FolD*, and *Fdh* suggests that members of *Wallacebacteriaceae* are not SAOB.

Based on the evidence found, it could be hypothesized that members of the family *Darwinibacteriaceae* are potential SAOBs, as they encode for the necessary genes to perform the reversed WL pathway coupled to the GCS. Within the intricate anaerobic digestion metabolic web, this pathway would allow these bacteria to perform the oxidation of acetate to H_2_ and CO_2_, reaction that would constitute a key step in AD, as hydrogenotrophic archaea would take this H_2_ as a substrate to produce methane. However, the family *Wallacebacteriaceae* lacks several key enzymatic of this pathway. Considering the relatively low abundance of this taxon in the analyzed bioreactors (as detailed in Supplementary Table 6), our hypothesis suggests that *Wallacebacteriaceae* may not play a significant role in the functioning of anaerobic digesters, unlike *Darwinibacteriaceae*. This could be attributed to the absence of a symbiotic relationship between *Wallacebacteriaceae* and hydrogenotrophic archaea.

## 4. Conclusions

In this work, a cryptic bacterial group, previously referred as MBA03 in SILVA database, was described for the first time by recovering and analysing representative genomes belonging to this taxon. Phylogenetic analyses confirmed that MBA03 is a representative of a new taxonomic order with the proposed name *Darwinibacteriales* ord. nov. Two different families can be distinguished within this order: *Darwinibacteriaceae* fam. nov. (also known as DTU010 according to the GTDB taxonomy) and *Wallacebacteriaceae* fam. nov. (also known as DTU012).

Order *Darwinibacteriales,* particularly family *Darwinibacteriaceae,* is among the most abundant genera in anaerobic digesters, regardless of operational and chemical parameters such as feedstock and type of reactor. *Darwinibacteriales* is a very specific bacterial group, being anaerobic digestion its main ecological niche. Bacteria from this order are also present in soils, sediments, plants and animal gut microbiomes at very low abundances. These materials are often used as substrates to feed anaerobic digesters, acting as a reservoir and an inoculum that allows *Darwinibacteriales* to thrive when the AD process begins.

The presence of the reversed WL pathway coupled to the GCS in the family *Darwinibacteriaceae* and the co-ocurrence of *Darwinibacteriales* with several hydrogenotrophic archaea (Otto et al., 2023) reveals that they are potential SAOB, and thus that they may act as competitors for acetate with acetoclastic archaea, favouring hydrogenotrophic methanogenesis. Furthermore, these bacteria seem to be important during hydrolysis, since *Darwinibacteriales* harbour a wide repertoire of hydrolytic enzymes (beta-N-acetylhexosaminidase, pullulanase, beta-glucosidase, chitinase, alpha-amylase). Nevertheless, it is important to highlight that this description has been entirely based on genome characterization, so its culture and further laboratory analyses are needed to confirm the hypotheses made.

The high abundance of *Darwinibacteriales* in AD environments strongly suggests an exaptation mechanism to this industrial process. In other words, *Darwinibacteriales* are naturally occurring hydrolytic and potentially SAO bacteria that are pre-adapted to AD and are thus massively enriched in biogas producing and other AD facilities, where they may play an industrially relevant role that our results have only begun to reveal.

## Supplementary Methods

### Sample description

As part of the MICRO4BIOGAS project, a total of 80 samples were collected from anaerobic digestion systems at 45 large-scale reactors with different operational conditions and feedstocks. These samples, together with the corresponding metadata, were used to investigate the chemical and taxonomic profiles (Otto et al., 2023). The first 40 samples were collected by members of the Technische Universität Dresden (TUD, Dresden, Germany) from 25 different biogas plants in Germany and Austria, while the second set of 40 samples was collected by the team members from Bioclear Earth B.V. (Groningen, Netherlands) from 20 different biogas plants in the Netherlands. A subset of 30 of these samples was selected to be analysed through metagenomic sequencing. This subset included samples enriched in MBA03, hydrolytic, acidogenic and acetogenic bacteria and/or methanogenic archaea. A full description of the samples can be found in Supplementary Table 1.

### DNA extraction and sequencing

The sludge samples were washed before DNA extraction to reduce the number of inhibiting substances. Specifically, 5-10 mL of each sample was mixed with 40 mL of sterile phosphate-buffered saline (PBS). Subsequently, the samples were centrifuged at 20,000 g for 10 min and the supernatants were discarded. This process was repeated twice, and the final pellets were used for DNA extraction with the DNeasy® PowerSoil® Pro kit (QIAGEN, Germany) based on the manufacturer’s instructions, with an extra incubation step at 65 °C for 10 min after adding the C1 solution to increase the efficiency of cell lysis. Qubit x1 dsDNA HS Assay kit (Qubit 4.0 Fluorometer, Thermo Fisher) was used for DNA quantification. The metagenomic DNA was randomly fragmented by sonication and DNA fragments were end polished, A-tailed, and ligated with the full-length adapters for Illumina sequencing: 5’-AGA TCG GAA GAG CGT CGT GTA GGG AAA GAG TGT AGA TCT CGG TGG TCG CCG TAT CAT T -3’ (5’ Adapter), and 5’-GAT CGG AAG AGC ACA CGT CTG AAC TCC AGT CAC GGA TGA CTA TCT CGT ATG CCG TCT TCT GCT TG -3’ (3’ adapter). PCR amplification with P5 and indexed P7 oligos was carried out, and the PCR products were purified with the AMPure XP system. Then, the libraries were checked for size distribution by an Agilent 2100 Bioanalyzer (Agilent Technologies, CA, USA), and quantified by real-time PCR. Libraries were sequenced at Novogene (UK) using the NovaSeq 6000 Illumina platform (150 bp x 2). The average sequencing depth was 45M reads/sample (min.: 35M reads; max.: 55M reads).

### IMNGS filtering

IMNGS results were filtered with the objective of gathering all the samples by their sample type to see which were the potentially relevant ecological niches of MBA03. As tags describing ecological niches were very diverse and, in many cases, ambiguous, a series of in-house Python scripts were needed to carry out a reclassification of niche categories (https://github.com/roserpr/IMNGSanalysis.git).

To briefly explain the filtering process, samples containing the 16S rRNA sequence of interest in an abundance <0.1% were removed from the analyses. To reclassify the remaining samples, two subsequent strategies were followed: (1) description tags were filtered by the words “anaerobic”, “digester”, “biogas”, and “reactor”, assigning all of them to the same tag: “anaerobic digestion”; (2) the information of the remaining samples were scraped from NCBI. The output was manually curated to determine which samples belonged to “anaerobic digestion”, and other general tags were also reassigned to more specific classes (i.e., “metagenome” to “soil metagenome”) depending on the project description.

