## Supplementary Material for "Unveiling the ecology, taxonomy and metabolic capabilities of MBA03, a potential key player in anaerobic digestion"

**Description of *Darwinibacter* gen. nov.**

*Darwinibacter* gen. nov. (Dar.wi.ni.bac′ter, N.L. masc. n., *Darwinibacter* in honor of Charles Darwin, a naturalist, geologist and biologist, best known for his contributions to evolutionary biology, and bacter, ‘bacterium’). (This text designates the taxonomic rank (genus) and the etymology under SeqCode rules 26.4 and 26.5).

Members of this genus have been identified from large-scale anaerobic digester tanks in Germany, Austria and the Netherlands. The genomic analysis of the genomes suggest that members of this genus are chemoorganoheterotrophs that derive energy from anaerobic respiration through oxidative phosphorylation. They utilize inorganic molecules of ferric iron (Fe^3+^) as potential electron acceptors. These bacteria also possess sulphide:quinone oxidoreductase, an enzyme involved in the oxidation of sulphides, including hydrogen sulphide, enabling them to be chemolithotroph.

They have complete sets of genes for glycolysis, gluconeogenesis, and pyruvate oxidation pathways. Moreover, they contain genes encoding beta-N-acetylhexosaminidase for bacterial cell wall peptidoglycan degradation and recycling or pullulanase for starch degradation. *Darwinibacter* members also possess additional genes encoding chitinase for chitin degradation, and alpha amylase for starch hydrolysis. Members of *Darwinibacter* assimilate nitrogen by converting NH_3_ to organic nitrogen through the enzymatic activities of glutamine synthetase and glutamate (NADPH) synthase. Members of this genera are mobile, as indicated by the presence of flagella synthesis genes.

The nomenclatural type of the genus is *Darwinibacter acetoxidans* gen. nov., sp. nov.

**Description of *Darwinibacter acetoxidans* sp. nov.**

*Darwinibacter acetoxidans* gen. nov., sp. nov. (a.cet.o’xi.dans. L. neut. n. acetum, vinegar; N.L. v. oxido, to oxidize; from Gr. masc. adj. oxys, acid or sour and in combined words indicating oxygen; N.L. part. adj. acetoxidans, acetate-oxidizing).

The phylogenetically closest species, according to EzBiocloud (Yoon et al., 2017) was *Parageobacillus galactosidasius* (NR_108140.1), with a 94.4% completeness and 88.64% similarity values. At a phylogenomic level, *Anoxybacter fermentans* (GCF_003991135.1) was the closest known species, with an ANI value of 86.06%.

It has the same physiological and metabolic characteristics as those which define the genus.

The MAG named M4b_bin25, representing the nomenclature type, was obtained from a mesophilic plug flow fermenter in the Netherlands fed with industrial wastewater. The type genome sequence of this species is 2.166.877 base pairs, consists of 83 contigs and has a G + C content of 59.19%. Completeness and contamination are 95.73% and 3.67% respectively, as estimated with CheckM (v1.2.2). ANI comparisons between this genome and those of closely related species (class *Clostridia* and *Limnochordia*) were below 86%, supporting the delineation of this taxon as unique and distinct from other species in the genus (Supplementary Table 8).

**Description of *Darwinibacteriaceae* fam. nov.**

*Darwinibacteriaceae* fam. nov. (Dar.wi.ni. bac.te.ri.a.ce’ae. N.L. masc. n. *Darwinibacter*, type genus of the family; L. fem. pl. n. suff. -aceae, ending to denote a family; N.L. fem. pl. n. *Darwinibacteriaceae*, the family of the genus *Darwinibacter*).

As parent taxon for the previously undescribed genus *Darwinibacter*, the family *Darwinibacteriaceae* is proposed. The type genus is *Darwinibacter* gen. nov. The description is the same as for the genus *Darwinibacter*. The taxon was represented by 36 genomes from anaerobic digestion tanks in Germany, Austria and Netherlands. To expand the study, 41 additional MAGs assigned to the GTDB family DTU010 by GTDB-Tk (Chaumeil et al., 2022) from other studies (<https://biogasmicrobiome.env.dtu.dk/>, Dyksma et al., 2020) were added.

**Description of *Wallacebacter* gen. nov.**

*Wallacebacter* gen. nov. (Wa.lla.ce.bac′ter, N.L. masc. n., Wallacebacter in honor of Alfred Russel Wallace, a naturalist, geologist and biologist, best known for independent discovery of the principle of natural selection, and bacter, ‘bacterium’).

Members of this genus have been identified from large-scale anaerobic digester tanks in Germany, Austria and the Netherlands. *Wallacebacteriacae* have complete sets of genes for glycolysis, gluconeogenesis, and pyruvate oxidation pathways. It encodes for the set of genes necessary for the biosynthesis of most amino acids, such as histidine, arginine, lysine, serine, glycine, proline, valine, methionine, isoleucine, leucine and tryptophan, meaning that *Wallaceae* representatives are auxotrophic for a smaller set of amino acids (Supplementary Table 6). It encodes for pullulanase for starch degradation, and beta-glucosidase for cellobiose metabolism, and also oligogalacturonide lyase, allowing them to degrade peptin. They are mobile, as indicated by the presence of flagella synthesis genes.

The nomenclatural type of the genus is *Wallacebacter cryptica* gen. nov., sp. nov.

**Description of *Wallacebacter cryptica* sp. nov.**

*Wallacebacter cryptica* gen. nov., sp. nov. (cryp’ti.ca. Gr. masc. adj. kryptikos, hidden; N.L. cryptica). It has the same characteristics as the one that define the genus.

The phylogenetically closest species, according to EzBioCloud (Yoon et al., 2017) was *Moorella thermoacetica* (NZ_CP012370.1)*,* with a 100% completeness and 88.61% similarity values. At a phylogenomic level, *Clostridium autoethanogenum* (GCF_001484725.1) was the closest known species, with an ANI value of 82.18%.

A MAG named M4b_bin43 representing nomenclatural type of the species was recovered from metagenomic sequencing from a mesophilic continuous stirred-tank reactor from Germany, fed mainly with manure and agricultural crops. The type genome sequence of this species is 2.331.610 base pairs, consists of 116 contigs and has a G + C content of 59.09%. Completeness and contamination are 94.89% and 1.98% respectively, as estimated with CheckM (v1.2.2). ANI comparisons between this genome and those of closely related species (class *Clostridia* and *Limnochordia*) were below 86%, supporting the delineation of this taxon as unique and distinct from other species in the genus (Supplementary Table 8).

**Description of *Wallacebacteriaceae* fam. nov.**

*Wallacebacteriaceae* is proposed (Wa.lla.ce.bac.te.ri.a.ce’ae. N.L. masc. n. Wallacebacter, type genus of the family; L. fem. pl. n. suff. -aceae, ending to denote a family; N.L. fem. pl. n.

As parent taxon for the previously undescribed genus *Wallacebacter*, family *Wallacebacteriaceae* is proposed *Wallacebacteriacea*, the family of the genus *Wallacebacter*. The type genus is *Wallacebacter* gen. nov.

The description is the same as for the genus *Wallacebacter*. The taxon was represented by 9 genomes from anaerobic digestion tanks in Germany, Austria and Netherlands. To expand the study, 16 additional MAGs assigned to the GTDB family DTU012 by GTDB-Tk (Chaumeil et al., 2022) from other studies (<https://biogasmicrobiome.env.dtu.dk/>, Dyksma et al., 2020) were added.

**Description of *Darwinibacteriales* ord. nov.**

As parent taxon for the previously undescribed families *Darwinibacteriaceae* and *Wallacebacteriaceae, Darwinibacteriales* is proposed (ord. nov. (Dar.wi.ni. bac.te.ri.a´les . N.L. masc. n. *Darwinibacter*, type genus of the order; L. fem. pl. n. suff. -ales, ending to denote an order; N.L. fem. pl. n. *Darwinibacteriales*, the *Darwinibacter* order). The type genus is *Darwinibacter* gen. nov.

The order was delimited based on phylogenetic (based on 16S rRNA gene) and phylogenomic analyses.
